# A trail camera imagery dataset of contrasting shrub and open microsites within the Carrizo Plain National Monument, San Luis Obispo County, California

**DOI:** 10.1101/044933

**Authors:** Taylor J. Noble, Christopher J. Lortie, Michael Westphal, H. Scott Butterfield

**Affiliations:** Department of Biology, York University, Toronto, Ontario, Canada.; U.S. Bureau of Land Management, Central Coast Field Office, Marina, California, United States.; The Nature Conservancy, San Francisco, California, United States.

**Keywords:** San Joaquin Desert, camera trapping, trail cameras, San Joaquin Valley, Carrizo Plain National Monument, San Luis Obispo County, endangered species, blunt-nosed leopard lizard, San Joaquin kit fox, shrub species, facilitation, *Ephedra californica*

## Abstract

**Background:** Carrizo Plain National Monument is one of the largest remaining patches of San Joaquin Desert left within the Central Valley of California. It is home to many threatened and endangered species including the San Joaquin kit fox, blunt-nosed leopard lizard, and giant kangaroo rat. The dominant plant lifeform is shrubs. The species *Ephedra californica* comprises a major proportion of the community within this region and likely also provides key ecosystem services. We used motion sensor trail cameras to examine interactions between animals and these shrubs. This technology is a less invasive alternative to other animal surveying methods such as line transects, radio tracking, and spotlight surveys. Cameras were placed within the shrub understory and in the open (i.e. non-canopied) microhabitats at ground level to estimate animal activity.

**Findings:** Trail cameras were successful in detecting the presence of animal species at shrub and open microhabitats. A total of 20 cameras were deployed from April 1^st^, 2015 to July 5^th^, 2015 at paired shrub/open microsites at three locations along Elkhorn Road in Carrizo Plain National Monument (35.1914° N, 119.7929° W). Each independent site was approximately 1 km^2^. Over 440,000 pictures (both of animals and triggers from vegetation moving in the wind) were taken during this time. The trigger rate was very high on the medium sensitivity camera setting in this desert ecosystem, and the rates did not differ between shrub and open microsites. The raw data (.jpeg images) are publicly available for download from GigaDB.

**Conclusions:** Motion sensor trail cameras are an effective, non-invasive alternative survey method for collecting data on presence/absence of desert animals. We detected mammals, reptiles, birds, and also insects in 0.4% of the images. We also successfully detected the Federally-listed blunt-nosed leopard lizard. A more extensive array of cameras within the Carrizo Plain National Monument could thus be an effective tool to estimate the presence of this species along with the presence of other animals.

## Data Description

### Background and Purpose of Data Collection

San Joaquin Desert habitat remains at less than 5% of its historical area. This region has largely been converted to irrigated agriculture and urban land uses (Germano *et al*. 2011, U.S. Fish and Wildlife Service 1998). These ecosystems are essential to the recovery and survival of a suite of endangered, threatened, and sensitive species including the endangered San Joaquin kit fox, giant kangaroo rat, and blunt-nosed leopard lizard (Germano *et al*. 2011, Germano *et al*. 2009, Prugh *et al*. 2010). Consequently, surveying animals within these remnant habitats within the region is important for conservation and management.

Three main core remnant San Joaquin Desert ecosystems remain in California. Carrizo Plain National Monument (35.1914° N, 119.7929° W), located in Southeastern San Luis Obispo County, is the largest (Germano *et al*. 2011). Precipitation at the monument ranges from 15 cm in the southeast to 25 cm in the northwest (Hijmans *et al*. 2005). Tests of animal camera technology were done in the Elkhorn Plain within the Monument on an elevated plain separated from the main valley floor of the Carrizo Plain by the San Andreas Fault (Germano *et al*. 1994). The area has been heavily invaded by non-native annual grasses including the following species: *Bromus madritensis, Erodium cicutarium*, and *Hordeum murinum* (Schiffman 1994, Gurney *et al*. 2015). The dominant shrubs are mormon tea (*Ephedra californica*) and saltbush (*Atriplex polycarpa*) (Stout *et al*. 2013).

Shrubs typically facilitate other plant and animal species in deserts by providing shelter, refuge, and resources (Lortie *et al*. 2015, Filazzola *et al*. 2014). We surveyed animal species with motion sensor camera traps from April through early July 2015 with a paired shrub/open structure to determine whether this technology was appropriate for assessing animal presence in deserts including under shrubs. The spatial partitioning of deserts into shrub-open habitat classes is a common method used to study positive interactions in deserts (Pescador *et al*. 2014), but has not been applied to the study of shrub-animal interactions in ecology. Imagery data is becoming increasingly common as a form of evidence for ecologists and conservation biologists (Swanson *et al*. 2015). The advantage of using cameras (a non-invasive survey method) is that it does not require handling the animals (which in this case includes many species that are threatened or endangered) and reduces overall disturbance to the study area. Additionally, the imagery is an important form of advocacy and engagement through citizen science.

### Camera Deployment and Imagery Collected

Cameras were set at the following three sites: Site 1: 35.192253°, -119.711794°, Site 2: 35.161870°, -119.672897°, Site 3: 35.114876°, -119.620541°. Sites were at least 2 km apart. This distance was selected to ensure that small mammals and lizards sightings at one site on a given day were independent of sightings at other sites. For instance, the daily movement of blunt-nosed leopard lizards usually ranges from 65 to 110 meters but can reach up to 300 m (Bailey *et al*. 2015). A total of 14 Primos trail cameras (Primos Hunting 2014) and 6 Reconyx trail cameras (Reconyx 2010) were deployed from April 1st through July 5^th^. Both models record at least 3 megapixels (with the ability to be set higher), with 2 sensitivity settings (low and high), replaceable batteries, IR camera, and both a motion sensor and IR sensor. Cameras were deployed in a paired design with one camera placed facing the north side of a shrub and the other facing away from the shrub at an open area at least 5 m away. Cameras were attached to 20 cm pegs anchored firmly in the ground. Vegetation was left intact and disturbance was minimized. From April until mid-May, cameras were set at each shrub/open pair from sunrise to sunset, and then moved to a new shrub/open pair the next day. From mid-May to July cameras were set at shrub open pairs for four days before being moved to a new shrub open pair to more intensively sample microhabitats at peak animal activity.

Sites were surveyed on consecutive days in a fully randomized order. At each set of paired shrub/open sites shrub size distance to nearest 3 shrubs, annual plant abundance, annual plant species composition, and annual plant density were recorded. Camera settings were also recorded. Camera images were examined to determine the presence of animals. Over 440,000 images (.jpeg) were collected over the season. We analyzed 100,000 of the images (prior to data analysis) to ensure image quality, presence of animals, date and temperature stamp, and data integrity. Animals were detected in 0.4% of the reviewed images. Using this approach San Joaquin antelope squirrels, jackrabbits, coyotes, blunt-nosed leopard lizards, whiptail lizards, side-blotched lizards, loggerhead shrikes, and grasshopper and butterfly species were observed at the Carrizo Plain National Monument in 2015 (Fig. 1). Images from cameras placed under both shrubs and in open microsites were clear and readable and the camera resolution was sufficiently resolved under low light and other challenging conditions to discern animals during both day and night (Fig. 2). The mean file size for each JPEG image was 750 kb. Filenames were encoded using the location the cameras were set at, the camera number and the date the pictures were downloaded. There were no camera failures during the season even though the units were placed in desert ecosystems with high temperatures.

## Potential Uses

This trail camera protocol can be used to estimate animal species presence/absence in desert ecosystems, and it is also effective under shrub canopies. The animal/insect capture rate of 0.4% suggests that extensive temporal and spatial sampling is required, particularly if the target animal species is relatively rare. However, extended surveys are possible because the trail cameras can be deployed for a week or more with little to no maintenance (Primos Hunting 2014, Reconyx 2010). Pricing and ease of deployment support the capacity to build more extended arrays within a region (Primos Hunting 2014, Reconyx 2010). Imagery data can be useful in describing vegetation characteristics and short-term micro-environmental disturbances. Temperature is encoded into every image, and these data can be extracted and used to evaluate fine-scale microclimatological differences.

## Availability of supporting materials and data

### Data Availability

The imagery dataset is deposited in the *Gigascience Database* repository, including the image, camera, and study design metadata. Images (.jpeg) are organized into folders based on the specific deployment site and date. Basic information about the habitat at the site (shrub or open), the type of trail camera used at that site and its settings, and the location of the survey site are also provided.

## Authors’ contributions

The data were compiled by TJN. CJL and TJN designed the experiment and wrote the manuscript. TJN conducted the field experiment, prepared the images for uploading, analyzed the data. CJL conducted a secondary round of screening of a subsample of the imagery for publication. MW assisted with experimental design, provided local knowledge, animal identification within imagery, participated in field deployment, and co-edited the manuscript. SB assisted with experimental design, provided local knowledge and expertise on the study area, assisted in field deployment and specific site selection, and co-edited the manuscript.

## Acknowledgements

TJN received funding for this study through the Fieldwork Cost Fund and Research Cost Fund from the Faculty of Graduate Studies at York University. Additional Funding was available through an NSERC Grant to CJL. MW and SB provided funding through grants to CJL from the Bureau of Land Management and The Nature Conservancy, respectively. We would like to express our thanks to the Bureau of Land Management, Bakersfield and Central Coast Field Offices for logistical support. Additional thanks to Kathy Sharum, Johna Hurl and the Carrizo Plain National Monument.

---

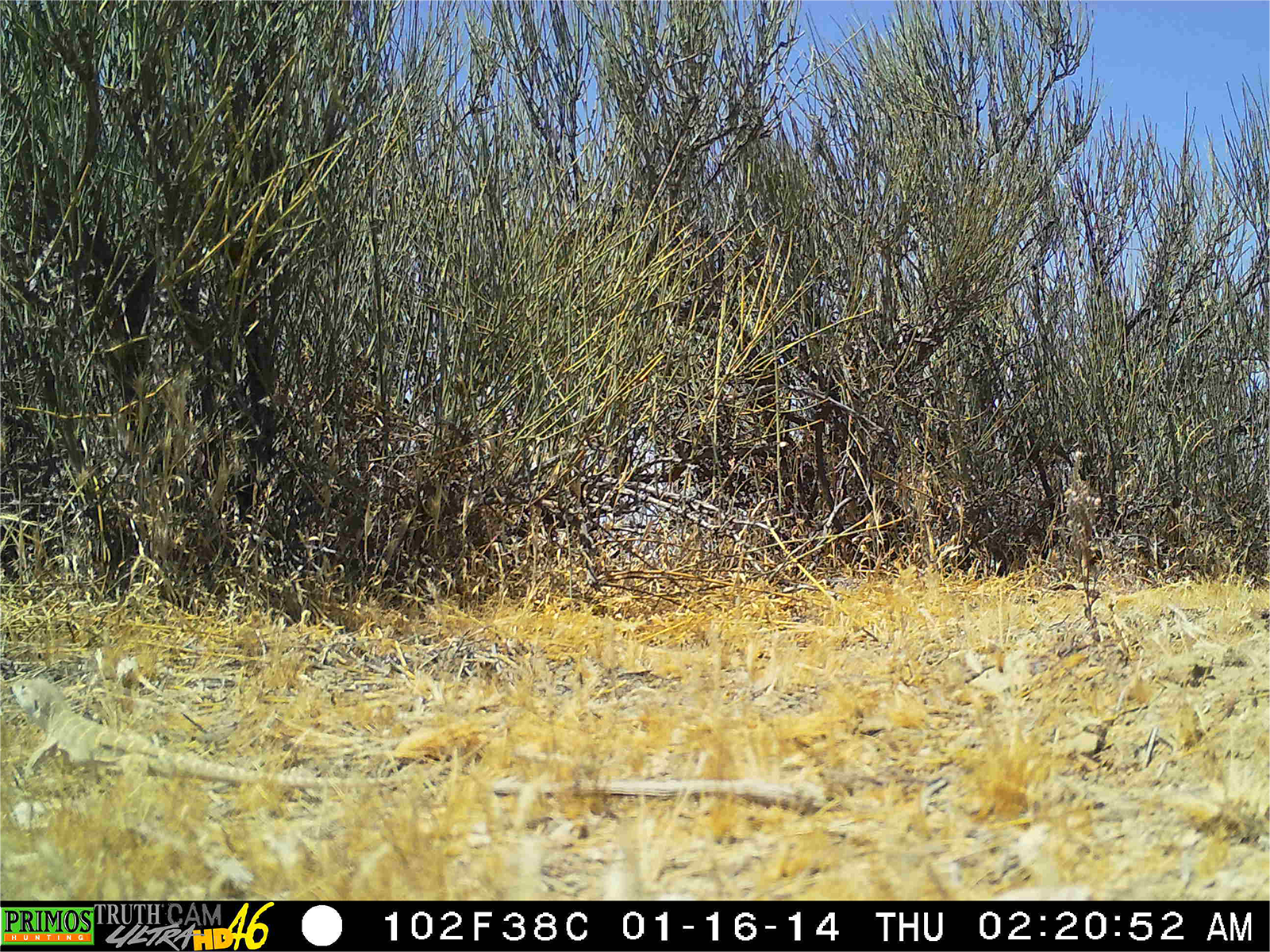

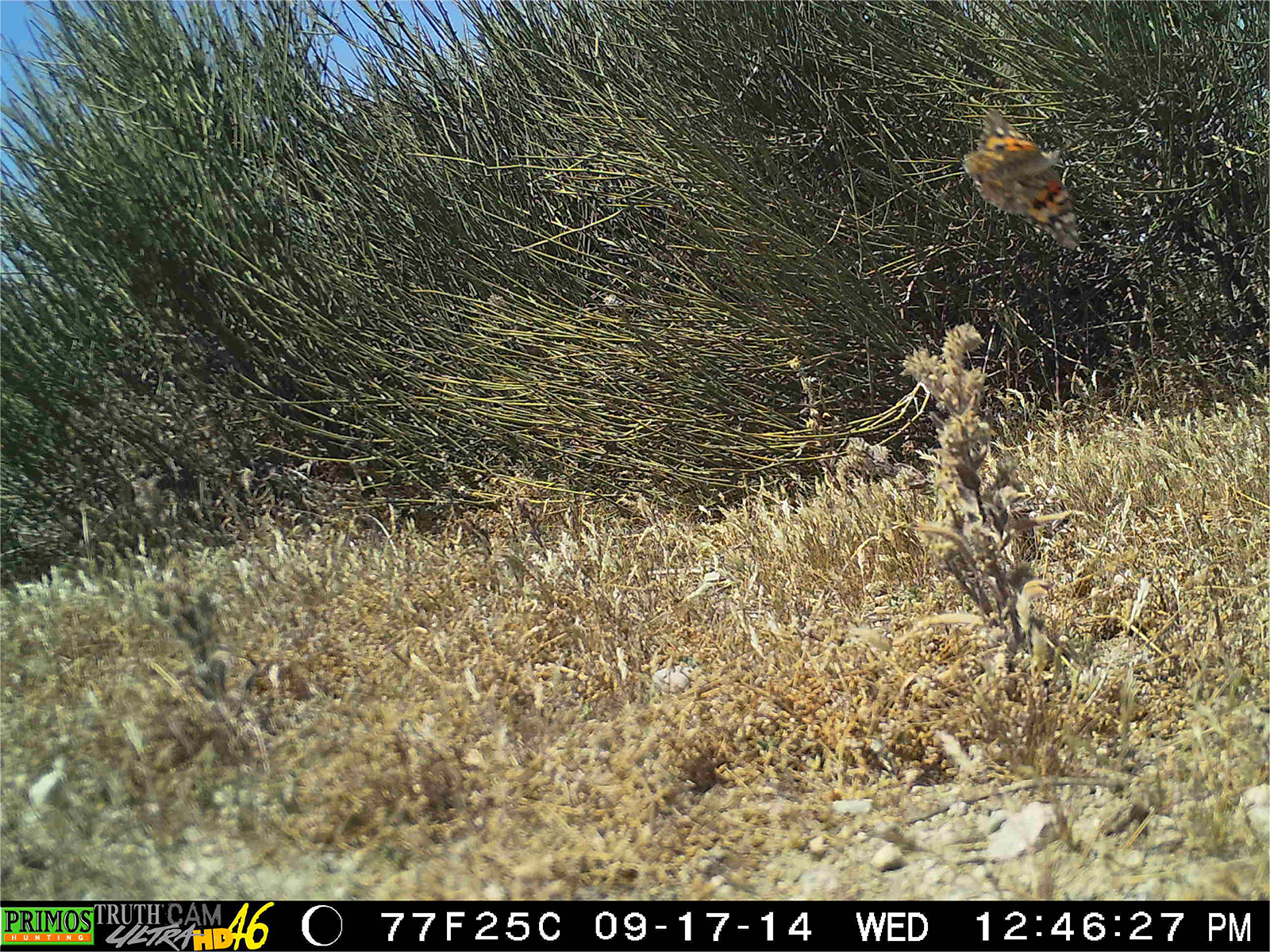

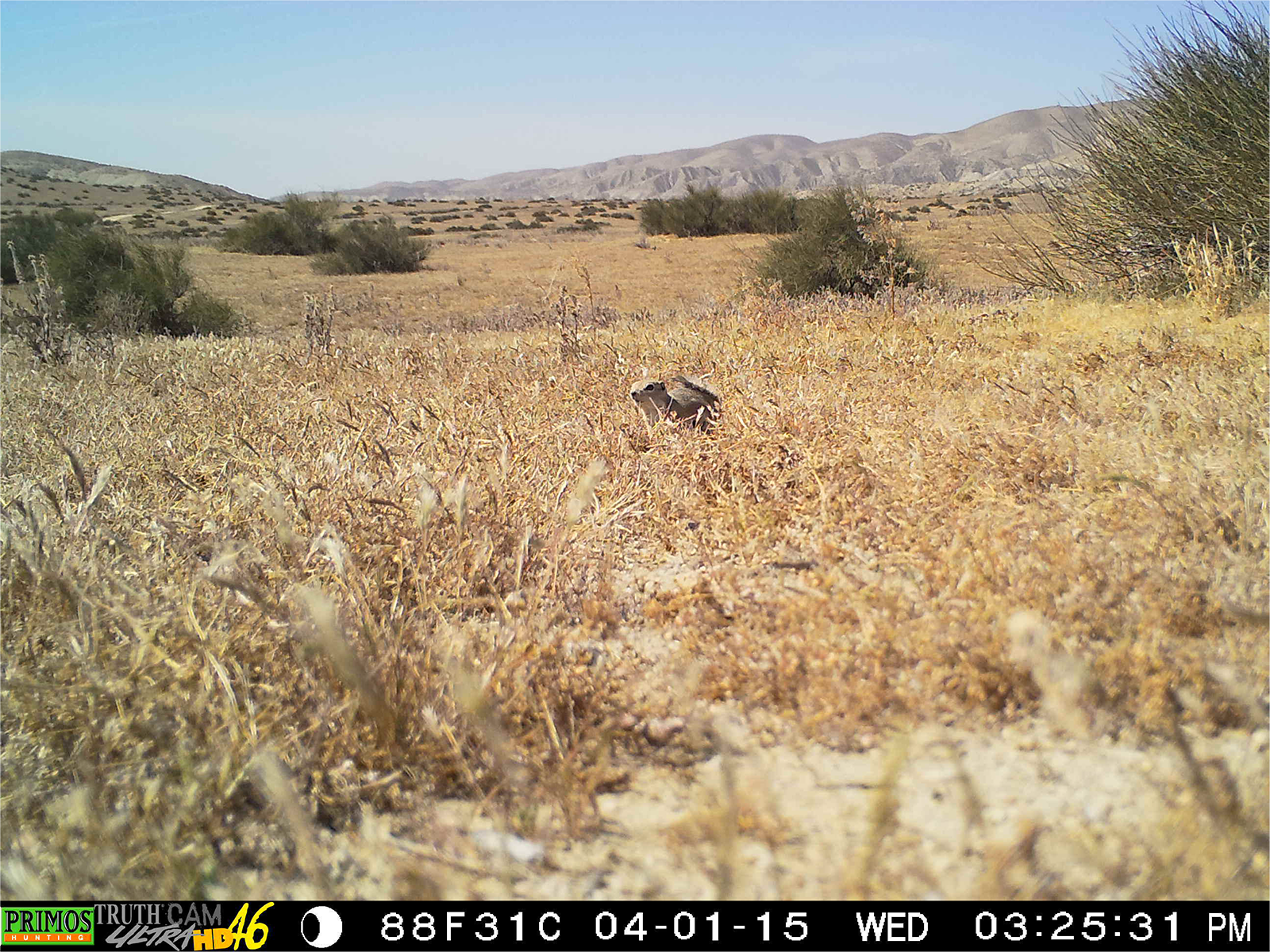

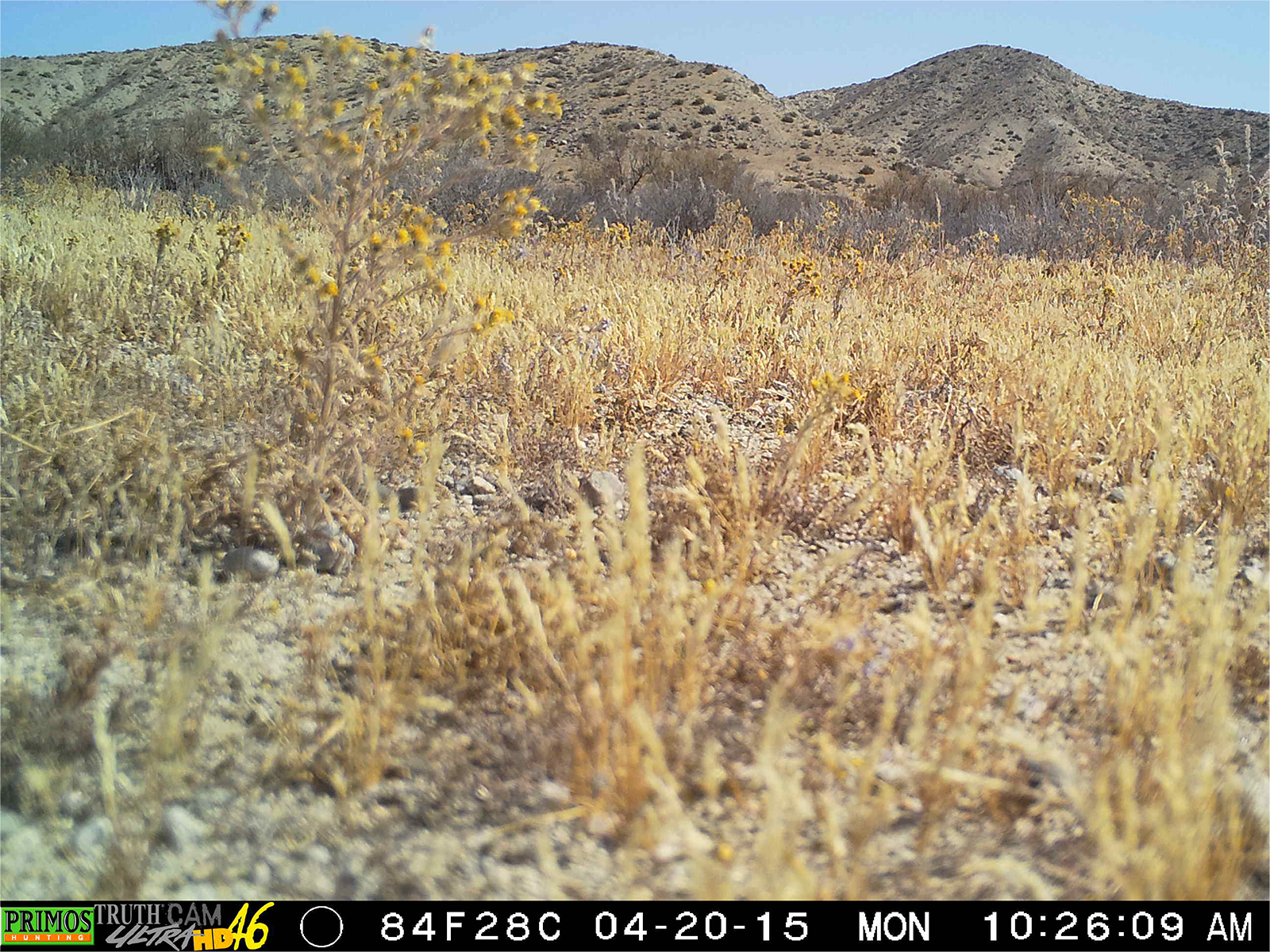

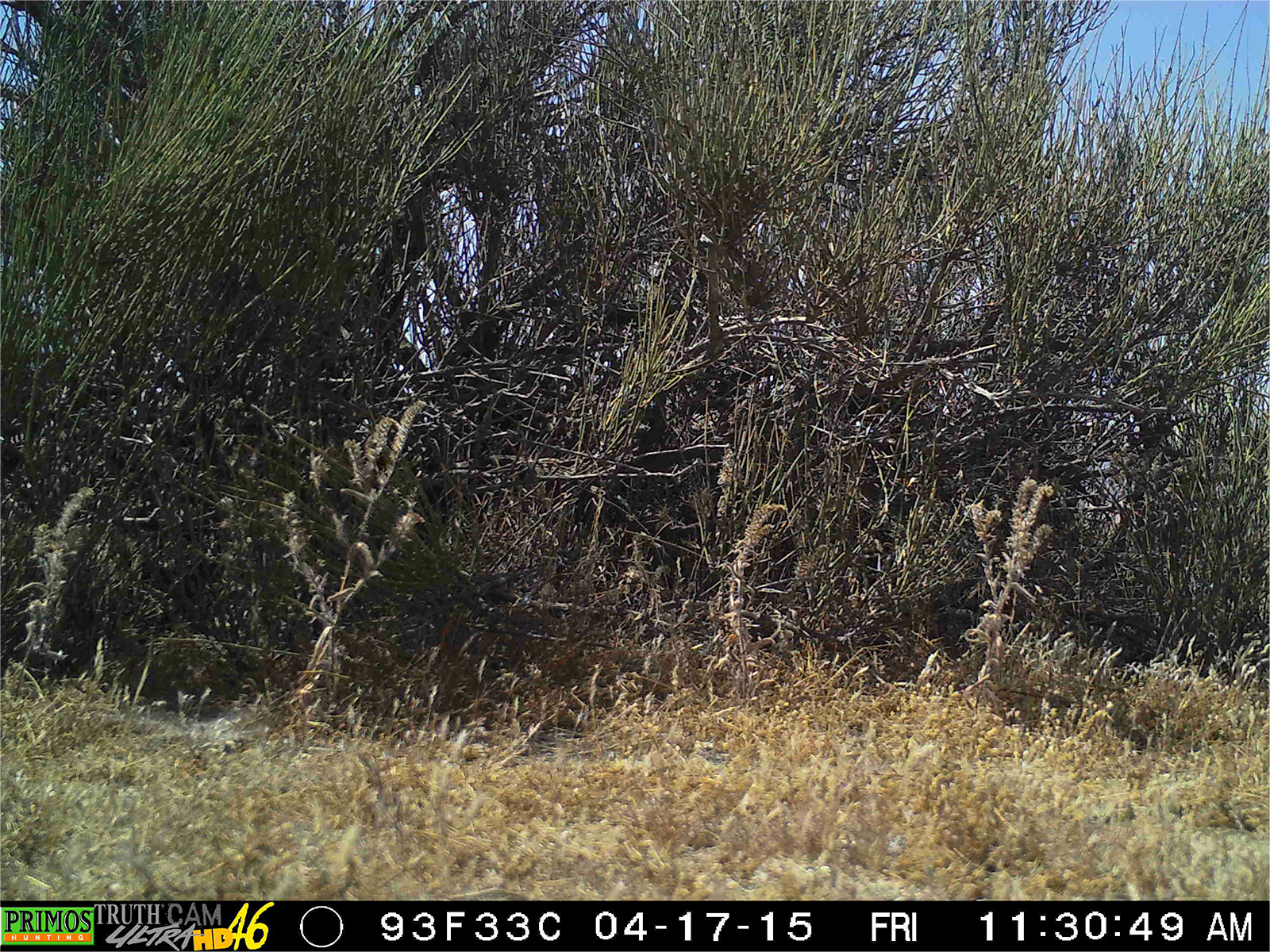

## References

Bailey C.V., Gemano D.J. 2015. Probability of occurrence of blunt-nosed leopard lizards on habitat patches of various sizes in the San Joaquin Desert of California. Western Wildlife. 2:23–28.

DeMay S.M., Rachlow J.L., Waits L.P., Becker P.A. 2015. Comparing telemetry and fecal DNA sampling methods to quantify survival and dispersal of juvenile pygmy rabbits. Wildlife Society Bulletin. 39(2):413–421.

Filazzola A., Lortie C.J. 2014. A systematic review and conceptual framework for the mechanistic pathways of nurse plants. Global Ecology and Biogeography 23: 1335–1345.

Germano D.J. 2009. The number of census days needed to detect blunt-nosed leopard lizards, Gambelia sila. California Fish and Game 95:106–109.

Germano D.J., Williams D.F., Tordoff III W. 1994. Effect of drought on blunt-nosed leopard lizards (Gambelia sila). Northwestern Naturalist 75:11–19.

Germano D.J., Rathbun G.B., Saslaw L.R., Cypher B.L., Cypher E.A., Vredenberg L. 2011. The San Joaquin Desert of California: Ecologically misunderstood and overlooked. Natural Areas Journal 31: 138–147.

Gurney C.M., Prugh L.R., Brashares J.S. 2015. Restoration of native plants is reduced by rodent-caused soil disturbance and seed removal. Rangeland Ecology and Management. 68(4):359–366.

Hijmans R.J., S.E. Cameron, J.L Parra, P.G. Jones and A. Jarvis, 2005. Very high resolution interpolated climate surfaces for global land areas. International Journal of Climatology 25: 1965-1978.

Lortie C.J., A. Filazzola, and D. Sotomayor. 2015. Functional assessment of animal interactions with shrub-facilitation complexes: a formal synthesis and conceptual framework. Functional Ecology. 30:41–51.

Pescador D.S., Chacon-Labella J., de la Cruz M., Escudero A. 2014. Maintaining distances with the engineer: patterns of coexistence in plant communities beyond the patch-bare dichotomy. New Phytologist. 204:140–148.

Primos Hunting. 2014. Primos Hunting Truthcam Ultra HD Instruction Manual. Available: http://www.primos.com/products/game-cameras/. Accessed Jan. 20th, 2016.

Prugh L. & Brashares J. (2010) Basking in the moonlight? Effect of illumination on capture success of the endangered giant kangaroo rat. Journal of Mammalogy, 91, 1205–1212.

Reconyx. 2010. Hyperfire High Performance Camera Instruction Manual. Available: http://www.reconyx.com/page/user-guides. Accessed Jan. 20th, 2016.

Schiffman P. M. 1994. Promotion of exotic weed establishment by endangered giant kangaroo rats (Dipodomys ingens) in a California grassland. Biodiversity and Conservation 3:524–537.

Stout D., Buck-Diaz J., Taylor S. & Evens J. (2013) Vegetation mapping and accuracy assessment report for Carrizo Plain National Monument. California Native Plants Society. https://www.cnps.org/cnps/vegetation/pdf/carrizo-mapping_rpt2013.pdf. Accessed Jan. 20th, 2016.

Swanson A., Kosmala M., Lintott C., Simpson R., Smith A., Packer C. 2015. Snapshot Serengeti, high-frequency annotated camera trap images of 40 mammalian species in an African savanna. Scientific Data. DOI: 10.1038

U.S. Fish and Wildlife Service. 1998. Recovery plan for upland species of the San Joaquin Valley, California. Portland, OR. 1-319.

